# Transitioning from global to local computational strategies during brain-machine interface learning

**DOI:** 10.1101/2020.03.05.978908

**Authors:** Nathaniel Bridges, Matthew Stickle, Karen Moxon

## Abstract

When learning to use a brain-machine interface (BMI), the brain modulates neuronal activity patterns, exploring and exploiting the state space defined by their neural manifold. Neurons directly involved in BMI control can display marked changes in their firing patterns during BMI learning. However, whether these changes extend to neurons not directly involved in BMI control remains unclear. To clarify this issue, we studied BMI learning in animals that were required to control the position of a platform with their neural signals. Animals that learned to control the platform and improved their performance in the task shifted from a global strategy, where both direct and indirect neurons modified their firing patterns, to a local strategy, where only direct neurons modified their firing rate, as animals became expert in the task. These results provide important insights into what differentiates successful and unsuccessful BMI learning and the computational mechanisms adopted by the neurons.

## Introduction

Since the first brain-machine interface (BMI) study in rodents (Chapin et al., 1999), it has been evident that animals can learn to modulate their neural activity to improve BMI control. To better understand this learning, researchers took advantage of high-density recording arrays, used a subset of neurons to control the BMI (direct neurons) and observed a separate population not used by the BMI (indirect neurons) (Ganguly et al., 2011) in order to understand the strategies used by the brain to acquire new learning. It has been proposed that there are two potential strategies the brain could use to learn new motor tasks. The first is a global strategy that suggests learning broadly engages the motor cortex and would be evidenced by both direct and indirect neurons engaging in the task (Chase et al., 2012; Jarosiewicz et al., 2008). The second is a local strategy wherein only the subset of neurons required to gain reward would be engaged in the task and other neurons would not engage (Zhou et al., 2019).

However, data from these experiments are mixed. In some cases, researchers did not find any changes in neural firing patters of indirect neurons (Law et al., 2014), while others demonstrated changes largely restricted to direct neurons (Arduin et al., 2013; Arduin et al., 2014; Clancy et al., 2014), especially after the animals learned the task well (Clancy et al., 2014). In studies that showed changes in neural firing patterns of indirect neurons, a broad array of changes were reported including changes in overall firing rate (Fetz and Baker, 1973; Ganguly et al., 2011; Gulati et al., 2014), which could be dependent on reward-timing (Hira et al., 2014), preferred direction (Ganguly et al., 2011; Hwang et al., 2013), latency or even coherency with slow-wave activity (Gulati et al., 2014).

One possibility is that the changes in neuronal firing patters in response to learning is dependent on when, during the learning process, the neuronal activity was assessed. Most BMI tasks require the animal to undergo some operant conditioning even before electrode arrays are implanted and this pre-training could impact BMI training. It is possible that prior conditioning alters the likelihood of observing changes in indirect neurons. Moreover, prior conditioning pre-selects animals that could learn the task, eliminating the ability to compare changes between animals that learned to those that did not, an important control group.

To overcome these limitations of pre-training, we developed a tilt BMI task (Figure 1A) that randomly applies different types of tilts to a platform while the animal works to maintain its center of mass task (Bridges et al., 2018). Neuronal activity is recorded and if the tilt type is correctly decoded by our classifier (Foffani and Moxon, 2004; Knudsen et al., 2014) the platform is returned to the neutral position, a natural reward. If the decoder incorrectly classifies the tilt type, the platform continues to tilt to an extreme position, a natural punishment. The task is relatively natural and does not require pre-training, making it possible to study BMI control in completely naïve animals. We used this task to study changes in performance of the BMI, which was determined by the ability of our classifier to discriminate between four different tilt types on a trial-by-trial basis. We studied the impact of learning, increases in performance, on the underlying cellular and network properties of direct and indirect neurons.

**Figure 1:**
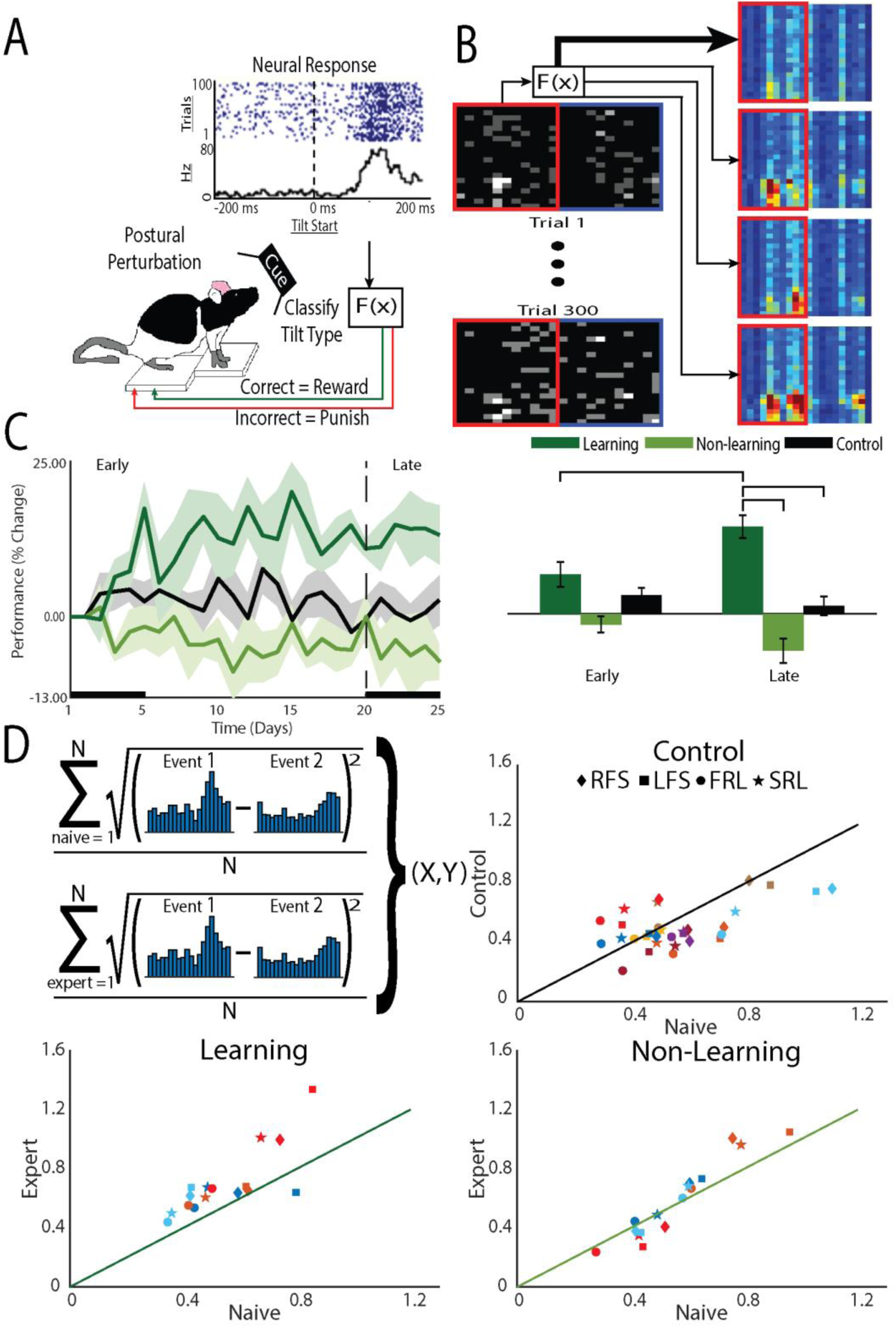
The BMI tilt task requires animals to learn to control a tilting platform. The task relies on the animal’s natural instincts and, therefore, does not require training to learn the task before the animal is expected to perform the BMI control task. (A) Animals stood on a platform that was tilted at a random speed in a random direction (clockwise or counterclockwise). (B) The PSTH-based classifier was used to decode the type of tilt from the neural activity and, if the tilt type was correctly classified, the platform was restored to the neutral position, a natural reward for the animal. If the tilt was incorrectly classified, the tilt continued to an extreme position, a natural punishment for the animal. (C) Performance of the animal in the BMI task was determined by the percentage of correctly classified tilts. Starting performance (Day 0) varied across animals so performance was normalized to Day 0 performance. Based on changes in their on-line performance over the five week period, animals were divided into learners, those that improved performance, and nonlearners, those that did not. At the conclusion of each recording session, off-line decoding was done using a leave-one out cross-validation on the recorded data to confirm the division into learners and nonlearners. (D) To assess the role of the decoder on learning, the difference in the PSTHs to different tilts was assessed and compared between early, when animals were naïve to the task and late when animals were expected to be expert at the task (upper left panel). For each animal (colors) for each tilt type (symbols: RFS, LFS, FRL, SRL) the difference in the PSTH during late (Expert) was plotted against the different early (Naïve). Points on the line demonstrate no differences between the PSTHs of experts compared to that of naive. Point above the line demonstrate differences greater difference between the PSTHs of experts compared to that of naïve. Greater differences between the PSTHs would improve decoder performance and indicate learning.

We hypothesized that animals would use a global strategy when learning to control the platform and, therefore, the learning mechanisms for direct and indirect neurons would be the same. While this was not what we observed, important differences between learners and nonlearners and direct and indirect neurons suggest that the brain uses a global strategy early to initiate learning but then switches to a local strategy, releasing indirect neurons from participation in conveying information about the task

## RESULTS

### Animals improve performance by increasing differences between neural responses

The task involves postural control and bilaterally engages the cortex (Bridges et al., 2018) (Deliagina et al., 2014; Jacobs and Horak, 2007). This allowed us to use one hemisphere for the direct neurons and the other hemisphere for the indirect neurons. Direct neurons were used control the platform as part of the BMI. The tilt was initiated and the neural activity within the first 200 ms was used to decode the type of tilt (out of four possible tilts, see Figure 1A). If the type of tilt was correctly classified, the animal was rewarded by having the platform return to its neutral position; incorrectly classified tilts resulted in a punishment and the platform continued to tilt to an extreme position. Indirect neurons were simultaneously recorded but not used in the decoder. Using separate hemispheres minimized interactions that might occur when using neurons from the same hemisphere, which are more likely to be anatomically or functionally linked and could potentially confound results. We examined changes in neuronal firing patterns as a function of performance in the task, as measured by the percentage of correctly classified tilts.

Importantly, in this task, there was no behavioral training before the animal was expected to perform the BMI control task (i.e. improve performance or the ability of the classifier to discriminate the correct tilt). Therefore, similar to the first experiments on operant conditioning of cortical unit activity (Fetz and Baker, 1973), this task did not require operant conditioning to train the animals in a behavior before they were placed into the BMI experiment. We compared performance of animals undergoing BMI training to a group of control animals subjected to the same tilts but without the ability for BMI control of the platform (control group). The task was difficult for the animals to learn with only about 50% learning BMI control creating two groups for comparison, learners and nonlearners. In addition, even for those animals that learned BMI control, the task took several days to achieve maximal performance creating two distinct learning phases: an early phase characterized by rapid, large performance improvements and a late phase characterized by long-term smaller performance improvements. We minimized decoder parameter changes across days of the experiment in an effort to maintain a stable cortical mapping to task outcomes (Ganguly and Carmena, 2009). The task also did not include any visual feedback and, therefore allowed a large state space within which the neuron could explore to determine how to improve their performance (Knudsen et al., 2012).

Because animals were learning to control the platform during each session, to examine what was learned, we examined changes in performance after building a decoder based on that day’s recording session, using a leave-one-out approach (off-line decoding). The decoder used the PSTH-based classifier (Foffani and Moxon, 2004) and classified the single trial neural responses (Figure 1B). Performance was defined as the proportion of correctly classified trials, normalized to Day 0 (first day on the tilt platform to collect data for decoder on Day 1). Off-line decoding performance supported the division into two group, learner and nonlearners (F(12,2)=7.1, p<0.01) but performance changed over time differently for the two groups [effect of phase: F(145,1)=0.2, p=0.69), interaction (F(145,2)=10.3, p<0.001]. Examining the learning curves Figure 1C) there was a noticeable early phase of learning, within the first 5-7 days, such that learners improved performance compared to their first, naive, day of BMI control (Day 1). There was also a late phase of learning. Learners became experts at controlling the platform characterized by an asymptotic leveling off of improvement in performance after day 15. The timing of these phases are similar to the early and late phase of learning demonstrated in humans (Wu et al., 2014) and BMI learning in non-human primates (NHPs) (Koralek et al., 2012).

Comparing group effects, as expected, only learning animals showed a greater change in performance in the late compared to the early phase (learning p<0.001; nonlearning p=0.08; control p=0.30). Within the early phase, there were no differences between groups, suggesting neurons recorded from nonlearners made attempts to improve performance but there was variability across animals. However, within the late phase, learners showed greater improvement in their performance compared to nonlearners (p<0.001) and control animals (p<0.01), while no differences existed between nonlearners and control animals (p=0.19). These data demonstrate that when animals learn to control the BMI, they continue to increase performance through the late phase (Zhou et al., 2019).

To understand why some animals were learners and some non-learners, we examined how neurons responded to the decoder and found that the neural activity of learners changed to enhance the performance of the decoder (Figure 1D). The on-line decoder used the PSTH-based classifier (Foffani and Moxon, 2004). This classifier uses the single neuron’s natural response to the tilt to drive the decoder. In practice, if the individual neuronal PSTHs to different tilts became more different from each other, then classifier performance would improve. For animals that learned, the difference between PSTHs for different tilt types was greater after they became experts at controlling the platform compared to their naïve performance before BMI learning began. This was less true for control animals and there was little to no change for non-learners (Figure 1D). Therefore, the choice of decoder will dictate, at least in part, the response of neurons to learning BMI control.

Furthermore, within a recording session, the distribution of incorrect trials was uniform across trials such that the cumulative sum of correct trials was linear (r^2^ > 0.99). Therefore, it is not as though improvement happens early or late within the recording session. This is different from improvements made by NHPs that are working in BMI tasks for which there was previous operant conditioning (Athalye et al., 2017). Here, improvement can be observed by the changes in slope of the cumulative performance improvement curves across weeks (Figure S1). The slope increased from early to late (F(1.76) = 3.142, p = 0.080) and was greater for learners compared to non-learners (F(1,76) = 5.729, p = 0.019) with a significant interaction (F(1,76) = 11.884, p = 0.001). In fact, for learners, the slope of this line increased as the animals learned the task, closely following performance (p<0.001) (Supplemental information Figure S1) but the slope of the line did not change for non-learners (p=0.21). Therefore, for learners, changes in firing patterns that matched the needs of the decoder occurred over the course of days to support learning.

### Brain strategies to learn BMI control

To understand the cellular computational mechanisms underlying learning, we examined how learning affected neuronal response patterns for the two types of neurons (i.e. direct or indirect) within learning groups y the early and late phases of learning (Figure 2). We first examined the effect of BMI learning on the change in information carried by individual neurons during learning. For learners, there was an effect of neuron type [F(7.7, 1)=12.46, p<0.01], with no effect of phase [F(2491,1)=0.73, p=0.39] but a significant interaction [F(2493,2)=6.5, p<0.005]. In the early phase, information increased more for both direct (p<0.05) and indirect (p<0.05) neurons compared to control, but they were not different from each other (p=0.833). Alternatively, in the late phase, the information for both direct (p<0.001) and indirect (p<0.05) neurons remained different from control but the change in direct was greater than that of indirect (p<0.1). In fact, only direct neurons increased information from early to late phase (p<0.001). Indirect neurons maintained their moderate increase in information in the late phase that they gained in the early phase with no additional increase (p=0.65). For non-learners, there was also an effect of neuron type [F(25.3,2)=3.46, p<0.05], with no effect of phase [F(2367,1)=1.7, p=0.19] and no interaction [F(2366,2)=0.004, p=1.0], suggesting that even if animals do not improve performance in the task, direct and indirect neurons are differentially affected by BMI training, albeit with smaller differences than that identified for learners. Therefore, early, the brain adopts a global strategy to improve BMI control but, overtime, transitions to using a local strategy such that only direct neurons continue to improve their encoding of the BMI control signal. This ability to transition from using a global to local strategy appears to be a property only found in learning animals.

**Figure 2:**
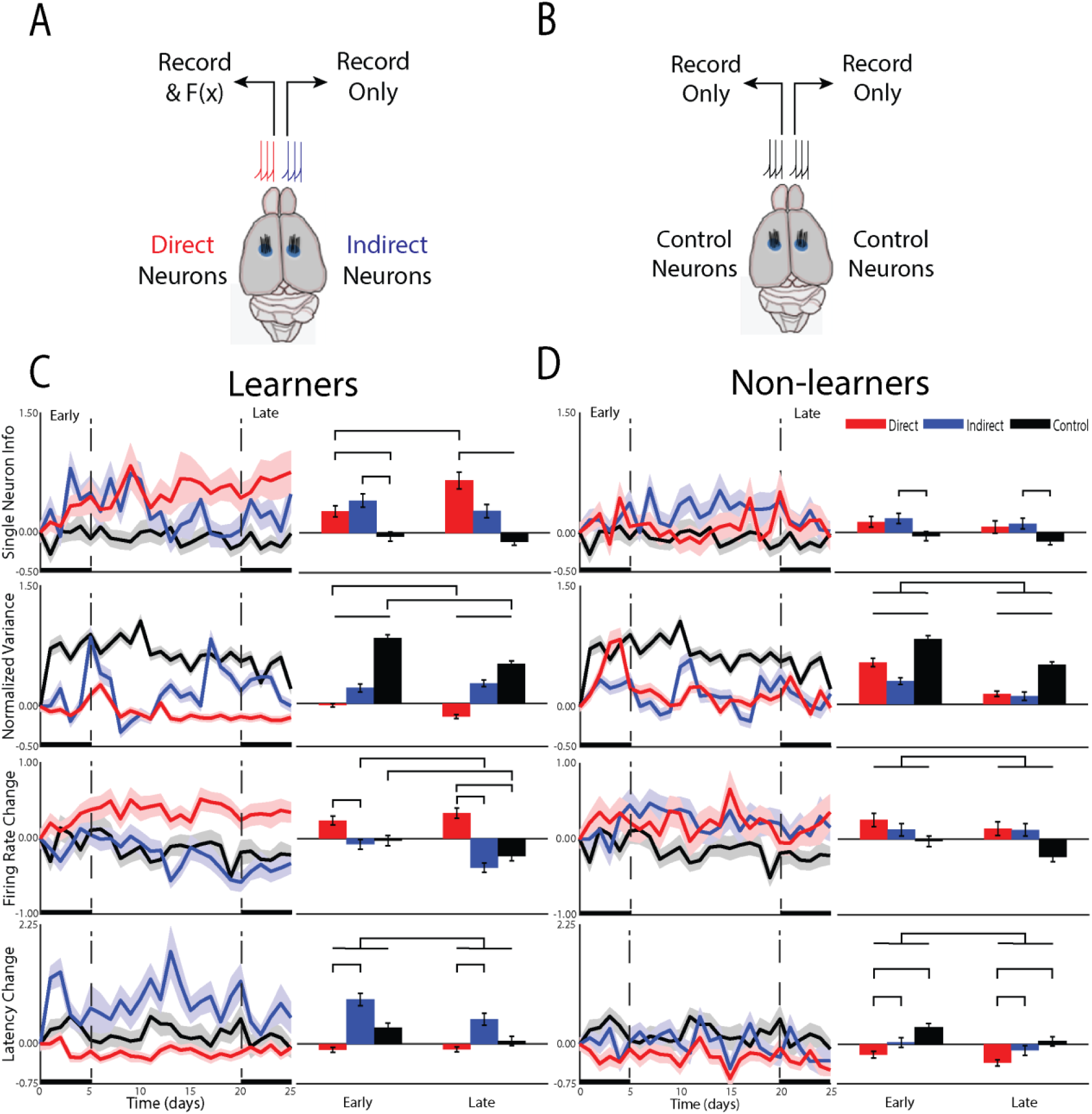
Differences in the cellular mechanisms underlying changes in the representation of information about the task. (A) For animals that underwent BMI, neurons from one hemisphere were used to drive the BMI (direct neurons) while neurons from the opposite hemisphere were recorded but not used in the BM (indirect). (B) a group of control animals had activity recorded from both hemispheres and their tilt types, rewards and punishments were yoked to one of the BMI animals but these animals did not undergo any BMI training. (C) Responses of single neurons compared between groups for animals that learned the task and (D) those that did not.

To understand this change in single neuron information, we examined changes in neural firing patterns. We first examined whether learning altered the neural variance (Churchland et al., 2006). Neural variance prior to the onset of movement has been shown to decrease— a measure of the preparedness of neurons to perform the task (Churchland et al., 2010). In animals that learned the BMI task, there was a significant effect of neuron type [F(2, 12,928))=229.30, p<0.001], phase [F(1,12,928)=17.2, p<0.001] and a significant interaction [F(2, 12928)=11.29, p<0.001]. Only direct neurons decreased their neural variance, continuing to reduce neural variance into the late phase compared to the early phase (p<0.001). In contrast, indirect neurons increased their neural variance early and with no additional increase late (p=0.38) but in a manner less than that of control animals who were tilted but had no ability to control the platform (p<0.001 for all comparisons between neuron type). For non-learners, there was an effect of neuron type [F(2,1264) =451.46, p<0.001] and phase [F(1, 1264)=66.18, p<0.001] with no interaction [F(2, 1264)= 2.21, p=0.11]. All neuron types increased their neural variance with less change from the early to late phase (p<0.01 for all comparisons). Again, neurons recorded from control animals had the largest increase, but this was followed by direct and then indirect. These data support previous studies that a reduction in neural variance is required for successful BMI control, further demonstrating that reducing neural variance is associated with learning the BMI task and is primarily reserved for direct neurons.

These differences in single neuron learning and neural variance between direct and indirect neuron types in the early phase of learning suggested that the brain was able to differentiate between direct and indirect neurons early on. To assess the underlying neuronal mechanisms that may contribute to this, we examined the effect of learning on the magnitude and latency of the neuronal response to platform tilts. For learners, changes in firing rate contributed to learning in the BMI task as there was an effect of neuron type [F(14.7,2)=24.3, p<0.001], phase [F(1912,1)=5.7, p<0.05] and a significant interaction [F(1913,2)=4.6, p<0.01]. In the early phase, while neither group was different from control, direct neurons increased their firing rate while indirect neurons decreased their firing rate and the difference between direct and indirect was significant (p<0.005) (Figure 2C). As the animals transitioned from the early to the late phase, direct neurons did not continue to increase their firing rate (early-late post hoc p=0.27) but indirect neurons continued to decrease their firing rate (p<0.01), similar to control (p<0.05). This effect on firing rate is intriguing and did not match changes in performance.

Learners also showed an effect of latency, but it was different from what was observed for firing rate. Overall, there was an effect of neuron type [F(13.5,2)=38.4, p<0.001] and an effect of phase [F(1911,1)=11.3, p=0.001] with no interaction [F(1911, 2)=2.49, p=0.08). This effect of phase was predominantly due to a larger increase in latency for indirect (p< 0.0001). The effect was attenuated in the late phase with the difference in latency between direct and indirect becoming less pronounced (Figure 2C).

For nonlearning animals, there were smaller changes in the firing rate and latency of the response. For firing rate, there was an overall reduction in the change during the late phase compared to the early [F(1604,1)=4.8, p<0.05] (Figure 2D). There was however, no difference between neuron types [F(15,2)=2.7, p=0.101] nor any interaction [F(1605,2)=0.342, p=0.710], suggesting that non-learning animals are unable to differentially modulate firing rate of direct or indirect neurons. For the latency of the response in nonlearning animals, there was an effect of phase [F(1603,1)=7.3, p<0.01) and an effect of group [F(14,2)=6.1, p<0.05] with no interaction [F(1604,2)=0.47, p=0.63). In fact, it is the direct neurons that are different from both indirect (p<0.005) and control (p<0.05) with no difference between indirect and control (p=0.24), suggesting that the direct neurons of nonlearners are working to improve performance but are unable to achieve an effect on the representation of information (Figure 2D). Therefore, as for learners, direct neurons of nonlearners were most affected by training in the BMI.

These data provide cellular computational mechanisms that support our informational results and suggest that learners initially use a global strategy with direct neurons changing their firing rate and indirect neurons changing their latency to learn to control the BMI. However, eventually the indirect neurons stop participating and, in the late phase, only the direct neurons contribute to learning BMI control. This is dependent on a continued reduction in neural variance with maintenance of firing rate increases and latency decreases while indirect neurons become less involved in the task. To understand the impact of these changes in firing rate and latency, we examined changes in information measures.

### Network Mechanism that Support BMI Learning

The differences in the neuronal responses of learners between direct and indirect neurons suggested network mechanisms underlying learning (Figure 3A,B). Specifically, the effects of firing rate on direct neurons and latency on indirect neurons suggested a differential role of temporal information. Temporal information is the information gained by considering the relative timing of spikes, or spike timing information, compared to the information conveyed by the overall number of spikes or spike count information (Foffani et al., 2004). The firing rate changes for direct neurons and latency changes for indirect neurons suggested a role for temporal information to be used by indirect neurons in the early phase of learning while indirect neurons relied on spike count information. In fact, the increase in information for indirect neurons relied on an increase in temporal information in the early phase (difference between spike count information and spiking timing information, p<0.05) and, as expected, this increase did not extend into the late phase (Figure 3A,B). For direct neurons, as expected, there was no added temporal information (p=0.13) in the early phase. However, in the late phase, the increase in information for direct neurons was through an increase in temporal information (p<0.05). Given that there was no further increase in firing rate nor change in latency, this increase in temporal information is most likely due to a reduction in jitter (Foffani et al., 2004; Foffani et al., 2007; Scaglione et al., 2014; Scaglione et al., 2011). Therefore, indirect neurons relied on increases in temporal information to increase information during the rapid, early improvement in performance while direct neurons improved temporal precision to gain further improvement in the late phase.

**Figure 3:**
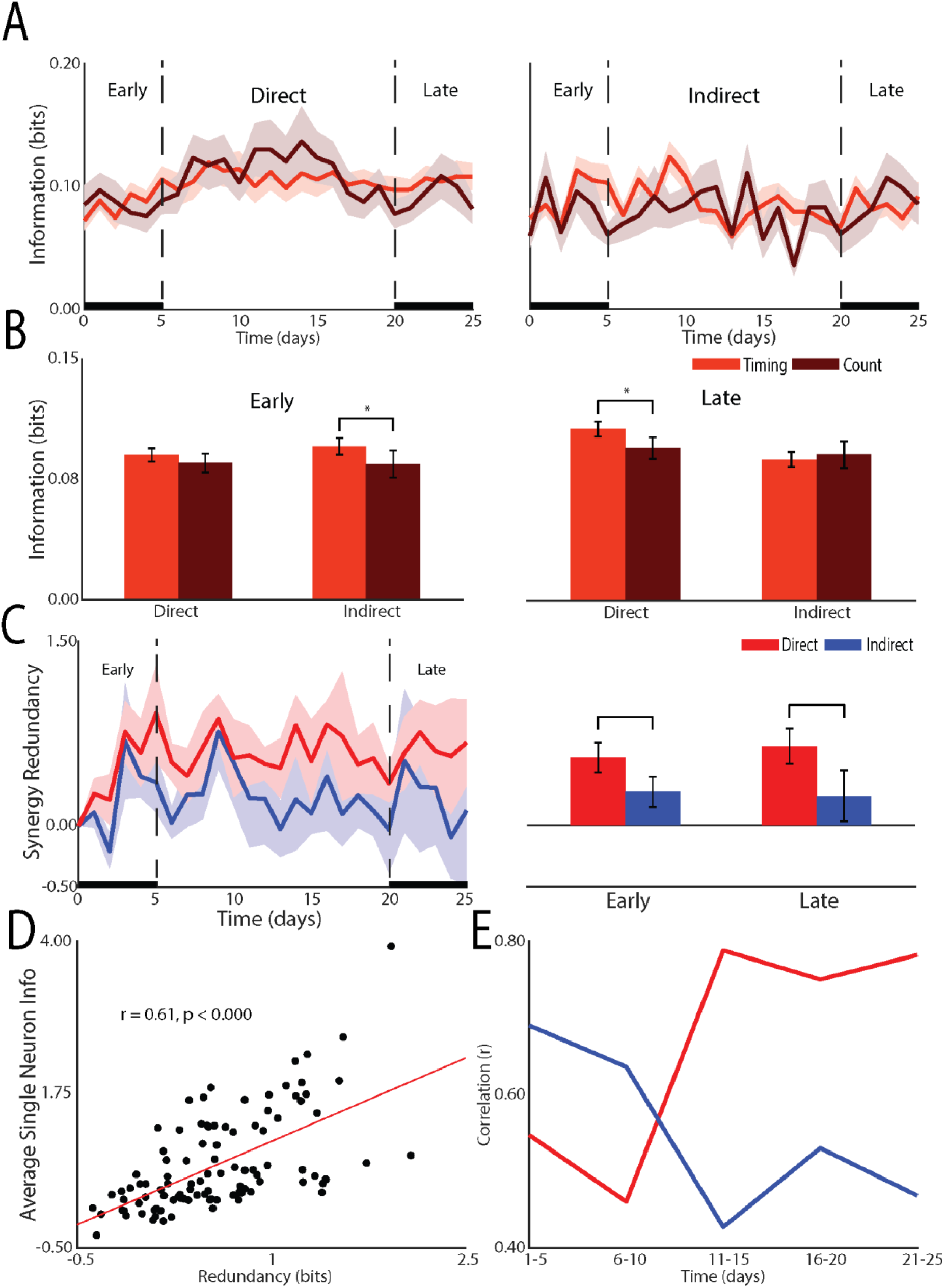
Effect of learning on network mechanisms. (A) Changes in the timing and count information for direct and indirect neurons across recording days. (B) Statistical comparison of changes in spike count and spike timing for during and indirect neurons druing the early (left panel) and late (right panel) phases. (C) Redundancy between population of direct and indirect neurons across resording days (left panel) and statistical comparisons for early and late (right panel. (D) Correlation between single neuron information and redundancy. (E)Changes in redundancy across recording days for direct and indirect neurons.

To gain a deeper understanding we examined the redundancy between neurons within the direct and indirect neurons (Figure 3C-E). As previously shown (So et al., 2012), single neuron information was well correlated to redundancy with the correlation increasing for direct neurons from early to late but decreasing for indirect neurons. In fact, the redundancy for direct neurons was greater than that of indirect neurons in both the early (p<0.01) and late (p<0.01) phase. Therefore, the transition from a global to a local strategy was accompanied by direct neurons transitioning from relying on spike count to relying on spiking timing information. This serves to increase the redundancy of the representation while indirect neurons are released from participating in the encoding of information about controlling the tilt platform.

## DISCUSSION

During the transition from the early to the late phase of learning, the brain changes from a global strategy—employing populations of neurons that directly influence BMI control and populations that do not influence BMI control—to a local strategy—optimizing for populations that directly influence BMI control while reducing irrelevant population activity. In the early phase, direct neurons reduced their neural variance and increased their firing rate to convey information through spike-count while the indirect neurons increased their latency to convey information through spike timing. As the brain shifted from a global to a local strategy, the direct neurons relied less on spike count and more on spiking timing, increasing the redundancy of the information representation. The indirect neurons decreased their latency and continued decreasing their firing rate, which further reduced their participation in conveying information about the tilt. The time course of local and global strategies and the reliance of different cellular computational mechanisms may explain the variability in research findings associated with indirect neuron investigations. In some cases, animals may rapidly transition from using a global to local strategy while in others the transition may not occur during the recording period (Chase et al., 2012). The nature of these changes is likely dependent on the parameters of the experiment and should be explored in future investigations.

Previous studies examining learning in a center-out reaching task recognized that a more global type of strategy was engaged early in learning (Chase et al., 2012; Zhou et al., 2019). In that case, the preferred direction of a subset of neurons used in the decoder were perturbed and the monkey had to relearn the mapping. In the early stage, both perturbed and unperturbed neurons as well as indirect neurons modulated their preferred direction, demonstrating a global strategy. It was suggested that the animal does this by re-aiming in the early phase of learning (Chase et al., 2012). In this study, we eliminated this type of approach (re-aiming) because the animal did not have continuous feedback during the trial regarding the successfulness of its attempt to control the platform. Further, because, in the reaching task, animals received operant training in the task prior to BMI control, learning peaked within the first recording session. In this study, learning was more difficult, allowing for a control group of nonleaners that demonstrate a role for single neuron learning and neural variance between neuron types. Specifically, the rapid changes in the early phase suggest that the brain has some understanding of direct neurons and indirect neurons in the early phase of learning.

In this study, we specifically defined a global strategy where both direct and indirect neurons participate in representing information about the task, rather than a behavioral response (re-aiming) that likely engages all neurons. A local strategy represented only direct neurons contributing to information about the task. Nevertheless, the reliance on cellular mechanisms early in learning suggest that cellular mechanisms are a universal property that is used to enhance performance regardless of whether the animal was previously trained via operant conditioning or whether the animal had continuous feedback regarding progress in the task. This may simplify the transition of BMI learning for restoration function although reexamining this after neurological injury or disease will be necessary.

### Local vs. Global Strategy

The ability to transition to a local strategy may be a key property of animals that successfully learn to use BMIs with training. This is supported by the fact that only learning animals were able to differentially modulate direct neuron information compared to indirect neurons when transitioning from the early to the late phase of learning. Non-learning animals continued to use a global strategy with changes in direct and indirect neuronal firing patterns correlated as training progressed. Therefore, a global strategy may be an exploratory step during which the brain attempts to identify the necessary neural transformation to gain reward. Since this is not necessarily the most efficient approach (Chase et al., 2012), once the transformation is identified, the brain adopts a local strategy such that cells not necessary for the transformation stop participating. These differences between learning and non-learning animal strategies may help explain why some animals learn and other do not, a well-known challenge in human BMI research known as the “BCI Illiteracy” problem.

Once the necessary transformation is identified, it would make sense to lock in the necessary local changes with structural neuroplastic changes, which take time to develop, in part because the brain must first identify the necessary local changes. Kleim and colleagues for example found that cortical synaptogenesis and motor map reorganization only occurred in the late phase of a motor-skill learning task in rats (Kleim et al., 2004). Global changes in contrast, may rely on more short-term functional neuromodulations. Since the learning in this task was solely dependent on reward rate, the animal could not adopt a behavioral strategy and the global strategy neurons adopted was an internal brain strategy.

In the earliest BMI studies in non-human primates (NHP), it was suggested that the animals internalize the robot arm (Carmena, 2013), consistent with early reports in human studies (Kennedy and Bakay, 1998). More recently, with the advent of improved technologies and more advanced software tools, BMI training has been used to gain insight into fundamental principles underlying learning. For example, changing the transfer function of the decoder such that it expects a different tuning curve of the cells (Jarosiewicz et al., 2008) showed that neurons will modify their tuning curve to gain rewards and that this process is reversible (Ganguly et al., 2009). More recently, the state space of the neural trajectories were assessed to determine if neurons would travel outside of their ‘natural’ state space, or manifold, if the decoder required it (Sadtler et al., 2014). It was difficult for neurons to do this and it has been suggested that there are intrinsic variables that limit the extent of change one could require of neurons (Hwang et al., 2013).

It has been suggested that learning takes place within existing motor repertoires underlying neural circuitry because learning is constrained to a low dimensional subspace of activity patterns or an intrinsic manifold (Sadtler et al., 2014). However, as shown here, learning takes place to solve the transformation of neural activity that gains reward (restoring the platform in this study). In many studies (Golub et al., 2018; Hwang et al., 2013; Sadtler et al., 2014), including the study presented here, the decoder defines how the neurons are required to change. The neurons simply work to improve the outcome of the classifier, which, in this study, did not require the neurons to move far from their more natural activity patterns. However, even when the required mapping was very different, for example, cells in the motor cortex working to control the tone of an auditory signal (Koralek et al., 2012), the change in neural firing patterns again matched the decoder. Therefore, it is unlikely that M1 uses a fixed repertoire of activity patterns but, rather, M1 learns a new mapping to acquire a new skill and this process creates a new low dimensional subspace of activity. The time it takes to learn a new mapping is dependent on the difference between the current mapping and the new mapping.

This study supports the idea that the brain undergoes an exploratory period, engaging a large number of neurons in the brain and trying different cellular and network strategies to gain reward. The direct neurons immediately identified changes in firing rate that allowed the PSTHs to different tilt types to move away from each, solving the transformation set by the decoder.

Changing firing rate does not allow the indirect neurons to participate in the transformation and a different strategy is attempted, changes in latency, leading to increased temporal information, but, again, the neurons do not participate in the transformation and eventually stop participating. This is made clear by the nonlearning control group. While it has been further suggested that learning on longer time scales (i.e. late phase) may involve adaptations outside of the manifold (Sadtler et al., 2014), our data do not support this prediction. Direct neurons simply lock in the transformation known to result in reward and do not change their relationship to the decoder. However, they do take advantage of additional mechanisms, including increasing temporal information that works to reduce the need for additional increases in firing rate while continuing to allow for increases in information.

It has been suggested that a local strategy requires the brain to solve the credit-assignment problem (Richards and Lillicrap, 2018) by identifying select neurons that contribute to desirable task outcomes. Solving the credit-assignment problem in BMI experiments is particularly challenging because the brain must parse out a relatively small subset of direct neurons. However, since we find differences between direct and indirect neuron types early, consistent with other studies (Zhou et al., 2019), the brain may have a built in mechanisms to solve the credit-assignment problem. Therefore, the early global strategy identified here is more than just simply identifying neurons that contribute to the desirable task outcome; this is known very early. Rather, neurons that can never contribute to the desired outcome continue to search for appropriate ways to contribute, for days in this case.

### Indirect/Direct Neurons Roles

It was initially suggested that a functional relationship exists between direct and indirect neurons, when neighboring direct and indirect neurons experienced similar modulation depths and response time changes (Gulati et al., 2014). However, in this experiment, direct and indirect neurons were separated across hemispheres, removing the likelihood of confounding factor of the close proximity of the cells. The differences noted here in firing rate and latency between direct and indirect neurons suggest that indirect neurons may contribute relevant information using independent mechanisms.

Latency differences seen in the direct neurons of this study are consistent with others. Manohar et al. found that direct neurons not only increased information but did so faster compared to neurons used in a manual control version of the task (Manohar et al., 2012). Further, Arduin et al. found that direct neurons respond faster than indirect neurons with BMI practice (Arduin et al., 2013), suggesting they act as “master” neurons by leading activation changes in cortical networks. Our firing rate differences parallel research showing that indirect neurons have smaller modulation depths compared to direct neurons (Law et al., 2014), which can decrease with time compared to manual control experimental conditions (Ganguly et al., 2009). Decreased firing rate might involve similar processes as those seen in neuroimaging studies where researchers identify decreased activation in experts compared to novices and/or as an individual learns (Babiloni et al., 2010; Guo et al., 2017; Yu and Yu, 2017) akin to the “neural efficiency” hypothesis. This may reflect a strategy employed by the brain to reduce the number of neurons involved in the task. BMI studies help to elucidate why this occurs; the brain starts by involving a large number of neurons and then down-selects those neurons that are not necessary to gain a desired outcome (i.e. righting of the platform).

Only learning animals showed a decrease in direct neuron neural variance as the animal transitioned from the early to late phase. This provides two important insights into skilled learning. First, neural variance may be used as a tool to probe the role of neuronal participation in a task. In the early phase, neural variance of direct neurons does not change much. In contrast, indirect neurons remain the same during the first few recording sessions but then begin to increase, reduce again and then increase, remaining relatively high during the late phase while that of direct neurons reduce. The reduction in neural variance before movement onset has been shown to be important for accurate task performance (Churchland et al., 2010; Churchland et al., 2006). The brain may use changes in neural variance to identify cells that are necessary for the task. Second, the reduction in neural variance, identified in well-trained animals, takes time to develop. These insights extend our understanding of the exploration of the neural state space while learning a task to include preparatory time before the onset of movement and that active modulation of indirect neurons in the early phase is important for learning.

### Redundancy and Neural Encoding

Interestingly, we found that redundancy increased for direct neurons yet decreased for indirect neurons as the animal transitioned from the early to late phase of BMI learning. Neuron ensembles most associated with BMI control display the highest information and redundancy. In one study, neurons from the hemisphere contralateral to the arm used to manually perform the task were used to train the decoder (direct neurons), while neurons in the ipsilateral hemisphere were recorded (indirect neurons) (So et al., 2012). In this case, direct neurons had more information and redundancy than indirect neurons as one might expect. Interestingly, when BMI control was switched such that the neurons from the ipsilateral hemisphere were used as direct neurons and the contralateral hemisphere for indirect, direct neurons (now in the ipsilateral cortex) still showed higher information and redundancy levels. Those data suggested that BMI related increases in information and redundancy are a property of BMI control. Our data confirm this result as only direct neurons increased redundancy with learning, even in the early phase when indirect neurons are increasing their representation about the task through latency changes but not contributing to BMI performance.

Additionally, computational work suggests that neural redundancy maximizes learning speed in motor cortical neurons (Takiyama and Okada, 2012). Our results are consistent with this idea in that direct neurons not only displayed more redundancy than indirect neurons but the redundancy was correlated with direct neural information. Interestingly, we found the degree of correlation between indirect neuron information and redundancy decreases as animals progress from early to late phase of BMI learning, while direct neuron correlational strength increases. This suggests that the brain takes an active role in increasing redundancy in neurons that contribute to task performance and reducing redundancy in those that do not as it transitions from using global to more local strategies. Various functions for neuronal redundancy have been suggested in rats (Narayanan et al., 2005) and primates (Reich et al., 2001) including that redundant neurons serve to safeguard against neural noise, which can be in the form of neurons misfiring (Puchalla et al., 2005) or unstable neural representations associated with motor learning (Rokni et al., 2007). Puchalla et al. (2005) proposed that redundancy allows for simpler high-order feature extraction via combinatorial codes in redundant systems. These theories of increased redundancy with BMI learning are consistent with Hebb’s hypothesis on cell assemblies (Hebb, 1949) that simply states while many neurons are part of the neuronal ensemble, on any given trial, only a subset need to fire in order to convey sufficient information about the task.

### Conclusion

In summary, these results resolve the myriad of neuronal response property changes for direct and indirect neurons identified for learning in a BMI by demonstrating that they are dependent on the phase of learning. Using an animal model that did not depend on prior operant conditioning was crucial to tease this apart as these mechanisms are dependent on the particular stage of learning. In fact, the early phase of rapid learning is defined by a global strategy where large number of neurons explore their neural space to identify mechanisms to contribute to earning a reward, in this case, righting the platform. In the late phase of lower rate of learning, indirect neurons stop participating in the task, while direct neurons adopt methods to improve temporal information and increase their redundancy in an effort to become more efficient. In general, the ability to shift from using global to more local strategies with time may be a key property of BMI learning.

## Materials and Methods

### Experimental Model and Subject Details

12 250-275 g female Long Evans rats, which were singly-housed under 12h light/dark cycle conditions, were used for this study. All animals were stereotaxically implanted bilaterally in the hindlimb sensorimotor cortex (Leergaard et al., 2004) within the infragranular layer (1.3 −1.5 mm) using 4 x 4 50 µm Teflon-insulated stainless steel microwires (MicroProbes for Life Sciences, USA). All surgical procedures were performed under general anesthesia (2-3% isoflurane in O_2_) via orotracheal intubation and pain was managed using Buprenorphine SR™ LAB (0.5 mg/kg; Wildlife Pharmaceuticals Inc., USA). Animals were allowed a week to recover from surgery before additional experimental manipulations. All animal procedures were conducted in accordance with the Drexel University Institutional Animal Care and Use Committee-approved protocols and followed established National Institutes of Health guidelines.

### Electrophysiology

Voltage waveforms were acquired at 40 kHz using a Multichannel Acquisition Processor (MAP, Plexon Inc., USA and discriminated into single neuron units using principle component cluster analysis and visual identification (Sort Client, Plexon Inc., USA).

### Behavioral task

At the start of the experiment, animals were subjected to four tilt types using a custom-built tilt platform coupled to a high performance brushless AC servo motor (J0400-301-4-000, Applied Motion Products, USA). LabVIEW (2015, National Instruments, USA) was used to initiate these tilts with a randomized order and 2-3 second intertrial interval for a total of 400 trials (i.e. 100 each tilt type). These trials were used to determine the baseline neural response state to each tilt type before rats received BMI training and is depicted as “day 0” in the results section. Importantly, neural responses to tilt perturbations did not influence outcomes of the tilt platform during baseline tilt sessions. In contrast, neural responses influenced the probability of tilt platform rewards and punishments during BMI training (see (Bridges et al., 2018), which immediately proceeded baseline tilt sessions for a total of 25 consecutive days (excluding Saturdays). Animals were split between the BMI (n=8) and control (n=4) animal groups. The neuronal activity recorded from BMI animals influenced task punishment/reward outcomes while neuronal activity recorded from control animals did not. In BMI animals, one hemisphere was used for direct neurons while the other was used for indirect. In contrast, neither hemisphere of control animals influenced task outcomes and the number and order of reward and non-reward tilts were matched to the BMI group.

### Performance

Performance of the animal in the BMI task was determined by the percentage of correctly classified tilts using our PSTH-based classifier. Starting performance (Day 0) varied across animals so performance was normalized to Day 0 performance. Rat neural response were used to discriminate between the four tilt types on a trial-by-trial basis using the PSTH-based classifier (Foffani and Moxon, 2004; Knudsen et al., 2014). Single-neuron average peri-event responses were created with a 20 ms bin size and a time window spanning ± 200 ms from the start of tilt from the previous day’s recording and used as templates for classifying tilt types. During BMI testing, the single trial response of the neurons was compared to the average neural response (i.e. templates) for each tilt type. The single trial was classified as belonging to the tilt type with the smallest Euclidean distance between template and single trial. As described in Bridges et al., care was taken to keep neuron definitions as similar as possible across recording days to minimize changes to the overall neuronal ensemble serving as an input to the decoder (Ganguly and Carmena, 2009). Neurons that stopped firing (relative to the previous day’s recording), however, were not used in the decoder for that day. Similarly, newly firing neurons not present in the previous day’s recording were recorded and saved but not used by the decoder for that day. If the new neuron was still present the following day it was used. Real-time decoding was implemented using Matlab (version 2012b, The MathWorks, Inc.).

## Data Analysis

For analyses, rats in the BMI group were split into “learning” animals (n=4), defined as animals with 5 consecutive days of increased task performance relative to first day performance and “non-learning” animals (n=4), which are those that did not meet this criteria (Bridges et al., 2018).Since both hemispheres in control animals were indirect, they were combined and treated as a single hemisphere for the following analyses. Additionally, only the first 300 trials were used for all analyses to minimize task-related fatigue effects. “Offline performance” was calculated using leave-one-out cross validation and normalized as a change in accuracy from the first day of BMI training. All other normalized measures were reflected as z-scores changes from the most recent baseline tilt session.

### Mutual Information Measures

Mutual information was calculated and is formally defined as:

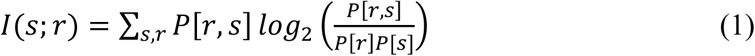

where *P[r], P[s]*, and *P[r,s]* correspond to the probability of the tilt-perturbation response *r*, the tilt perturbation stimulus *s* and their joint probability respectively. *I(s;r)*, which is measured in units of bits, was calculated for each neuron across the 25 days of the experiment using the actual and predicted tilt type confusion matrix generated when applying the classifier (Foffani et al., 2004). Residual bias for *I(s;r)* was then estimated using a bootstrapping procedure by pairing the trial responses and tilt types in a randomized order—effectively eliminating the association between the two (Magri et al., 2009). This procedure was performed 50 times, which provided an asymptotically stable estimate of random information within 0.001 bits, and the calculated bias was subtracted from *I(s;r)*. Only neurons with bias-corrected mutual information above 0 were used for information analyses. All information analyses were calculated using a 200 ms window from the start of tilt. Spike timing information was calculated using 20 ms bins, while spike count information was calculated using one 200 ms bin.

Redundancy was calculated and is formally defined as:

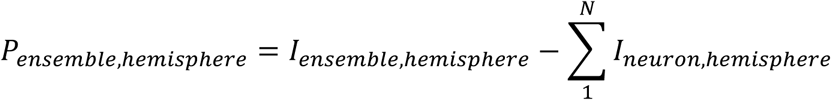

Where *I*_*ensemble,hemisphere*._corresponds to the mutual information calculated using all recorded neurons and *I*_*neuron,hemisphere*._corresponds to the mutual information of a single neuron. This analysis was performed separately for the direct and indirect neuron hemispheres. For plotting and conceptual purposes *P* measures were inverted such that increases in value corresponded to increases in redundancy. Pearson’s correlations were made between the average mutual information across neurons and redundancy z-score for each BMI training day per animal.

### Neural Response Property Measures

As previously described (Bridges et al., 2018), the following neural response measures were calculated: (a) Peak Response (PR): the PSTH bin with the maximum number of spikes divided by the total number of trials after subtracting the BFR. (b) Peak latency (PL): the time (i.e. bin) relative to tilt start the PR occurs. Only “responsive” neurons, defined as a neuron having at least 5 consecutive 2 ms above threshold PSTH bins and a response window significantly greater than the average background activity (one-sided paired t-test, p>.001), were used in this analysis. If a neuron was responsive to multiple tilt types, the most responsive case was used such that each neuron only contributed one observation to the analysis.

### Euclidian Distance

The Euclidian distances were calculated between tilt types in order to compare how an animal changed its performance and classification. The direct hemisphere from the learning and non-learning group were used while both indirect hemispheres from control were used as a reference. The Euclidian distance was found by taking each neuron’s PSTH for each tilt type from the naïve day, day 0, and the expert day, determined by performance. Each neuron’s PSTH was lined up bin-wise for two given tilts and the Euclidian distance was found according to the formula:

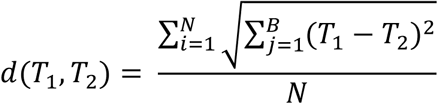

where T_1_ and T_2_ are the two PSTH tilt responses being compared and B represents the total bins. A bin size of 20ms from −200ms to 200ms was used to create each PSTH. The Euclidian distance was then averaged across neurons to find the population Euclidian distance and then plotted, where the x value was the naïve day distance and the y value was the expert day distance.

### Normalized Variance

The normalized variance was calculated (Churchland et al., 2006) and then z-scored for each of the four tilts. After determining that the z-scored normalized variance was not different between tilts, the z-scored normalized variance was averaged together to give a single value for each hemisphere for each day. The data was not smoothed as suggested, but the background spikes were binned in a single 200ms bin and was scaled by a 200ms constant and offset by an epsilon of 0.01 (Churchland et al., 2006).

### Intertrial Analysis

An intertrial analysis was conducted for each hemisphere for all animals to examine how the animal adapted to the new decoder each day. To do so, the cumulative sum was taken across trials, where a correct trial resulted with a +1 and an incorrect trial resulted with a +0. The cumulative sum data was then plotted and a linear line of best fit was found, along with a r^2^ value for each plot.

### Statistics

Statistical comparisons were made between the first (early phase) and last five days (late phase) of the experiment. A hierarchal linear modeling (HLM) approach was used for these comparisons to account for the fact that multiple neurons were recorded from the same animal (see (Bridges et al., 2018). This approach was used for all early to late phase comparisons except for spike timing and count differences (Figure 3), which used Mann Whitney U tests. We opted to use Mann Whitney U tests in this case because the HLM approach resulted in Hessian matrix issues, which was likely the result of redundant covariance parameters/overcomplicated model parameters. In all cases, Bonferroni-corrections were applied when appropriate. All error bars are in the form of standard error of the mean.

## Author Contributions

NB collected all of the data and NB and KM contributed to all other aspects of this paper, MS contributed to data analysis and figure production. This work was supported by grant NS096971 from the National Insitutes of Health and grant 1933751 from the National Science Foundation.

## Figure Legends

**Supplemental Figure 1:**
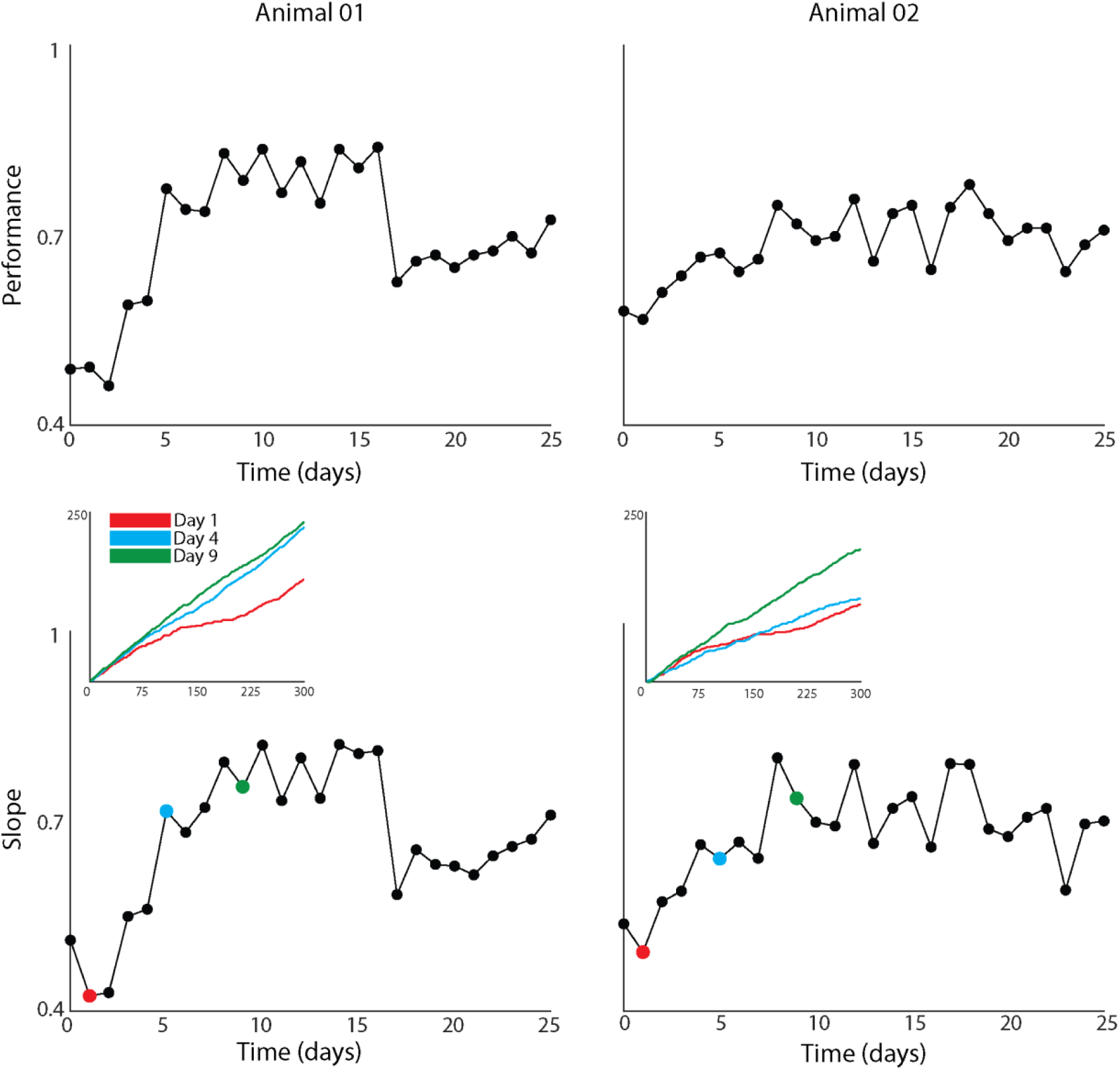
Examples showing the performance (top panels) and the slope of the cumulative sum of performance within a recording session (bottom panels) for two animals. Insets for bottom panels show the cumulative sum of the current trials during single recording sessions (Day 1, 4 and 9 as examples). From the insets, it is clear the cumulative sum of the correct trials is linear, also see text. The slope of the cumulative sum for each day a measure of how much was learned, the steeper the slope the more correct trials during that recording session. The slopes increase from earlier days (e.g. Day 1) to later days (e.g. Day 9). When the slope of the cumulative sum is plotted for each day (bottom panels) the shape of the curves are similar to that animal’s performance (top panels), suggesting that the rate at which animals learn during a session is well described by the overall performance of that session.

